# As48, a First-in-Class Dual-Function TREM2 Modulator: Receptor Activation and Shedding Inhibition

**DOI:** 10.1101/2025.08.23.671919

**Authors:** Sungwoo Cho, Farida El Gaamouch, Saurabh Upadhyay, Hossam Nada, Katarzyna Kuncewicz, Moustafa T. Gabr

## Abstract

Triggering receptor expressed on myeloid cells 2 (TREM2) dysfunction contributes to Alzheimer’s disease pathogenesis, yet current therapeutics cannot prevent ADAM-mediated receptor shedding that diminishes signaling efficacy. Using Affinity Selection-Mass Spectrometry (AS-MS) screening, we identified As48, a novel small molecule that binds TREM2 with high affinity. Biophysical validation confirmed s 7-fold selectivity over TREM1. Cellular assays demonstrated that As48 functions as a TREM2 agonist, activating SYK phosphorylation and enhancing microglial phagocytosis. Molecular docking and molecular dynamics simulations revealed that As48 binds near the cleavage region, establishing hydrogen bonds with Gly68 and reducing conformational flexibility in regions 58-102. Based on this structural insight, we investigated the effect of As48 on TREM2 ectodomain shedding and discovered inhibition of receptor shedding without affecting ADAM10/17 protease activities, representing the first small molecule with anti-shedding properties through conformational restriction of protease accessibility. Importantly, As48 displayed favorable pharmacokinetics with potential for brain permeability, supporting its translational relevance. Through its dual and simultaneous promotion of receptor activation and prevention of shedding, As48 represents a paradigm shift in TREM2 modulation and neuroinflammatory drug discovery.

**Abstract figure:** 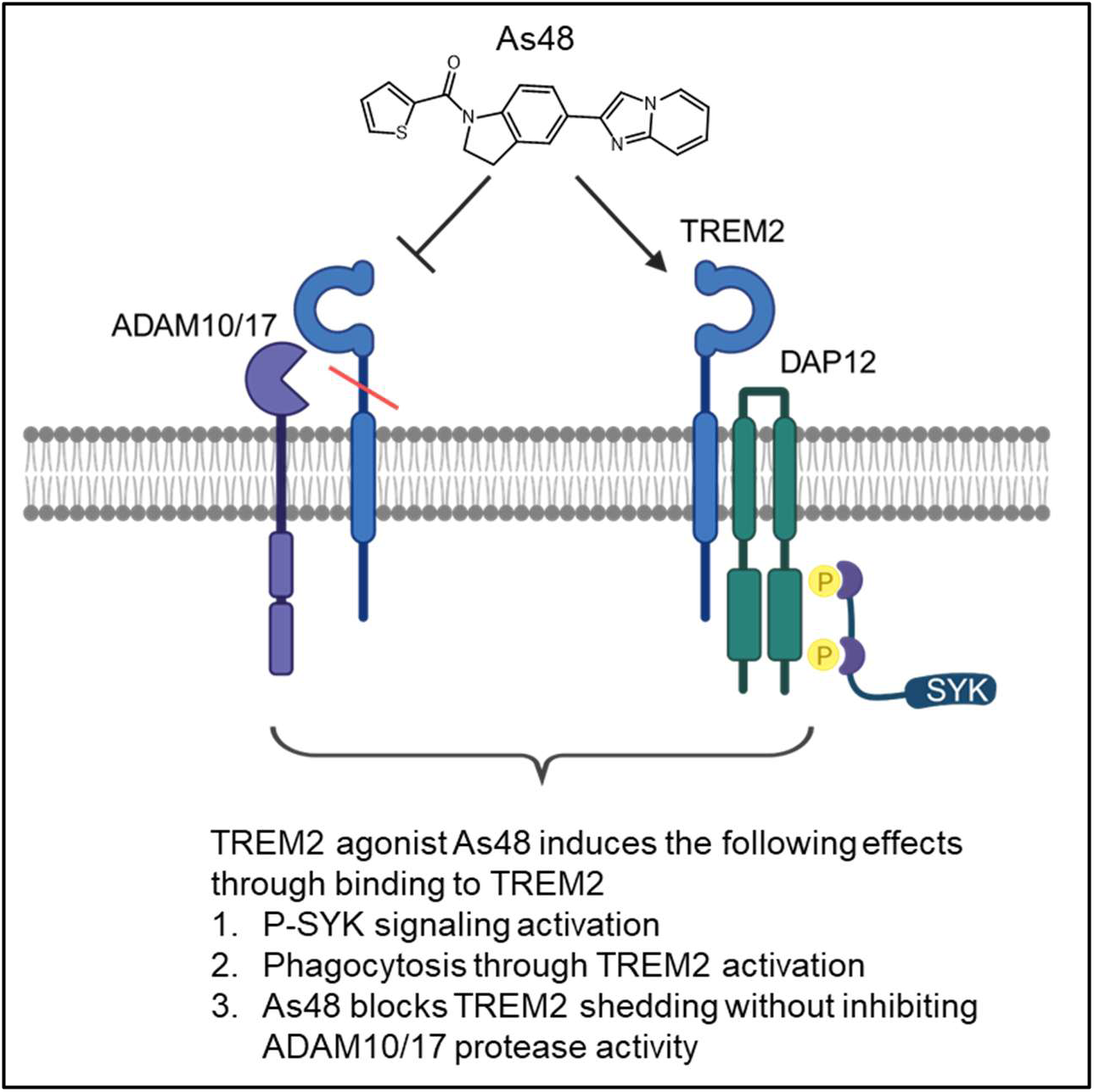

## Introduction

Triggering receptor expressed on myeloid cells 2 (TREM2) is a transmembrane immunoreceptor primarily expressed on microglia in the central nervous system (CNS) and tissue-resident macrophages in peripheral organs. This receptor plays a pivotal role in regulating microglial activation, phagocytosis, lipid metabolism, and inflammatory responses, making it a critical mediator of neuroinflammation and neurodegeneration (Colonna and Wang 2016, Deczkowska, Weiner et al. 2020). TREM2 recognizes lipid ligands and damage-associated molecular patterns, enabling microglia to sense and respond to pathological changes in the brain environment (Wang, Cella et al. 2015). TREM2 functions through association with the adaptor protein DAP12, initiating downstream signaling cascades that promote microglial survival, migration, and debris clearance (Wang, Cella et al. 2015, Kober and Brett 2017). Loss-of-function mutations in TREM2, including the well-characterized R47H and R62H variants, have been identified as significant risk factors for Alzheimer’s disease (AD), substantially increasing disease susceptibility by 2-5 fold (Guerreiro, Wojtas et al. 2013, Ulland and Colonna 2018). These mutations impair microglial function, leading to decreased amyloid-β (Aβ) plaque clearance, reduced lipid metabolism, and dysregulated inflammatory responses, ultimately contributing to disease pathogenesis (Griciuc, Serrano-Pozo et al. 2013, Keren-Shaul, Spinrad et al. 2017, Zhao, Wu et al. 2018). Recent single-cell RNA sequencing studies have identified disease-associated microglia (DAM) that depend on TREM2 signaling for their activation and neuroprotective functions (Keren-Shaul, Spinrad et al. 2017, Krasemann, Madore et al. 2017).

The critical role of TREM2 in neurodegeneration has spurred intensive efforts to develop therapeutic strategies targeting this receptor. Current approaches primarily focus on two modalities: monoclonal antibodies (mAbs) and small-molecule agonists (Fassler, Rappaport et al. 2021, Zhao, Xu et al. 2022, Mirescu 2024, Park, Han et al. 2024). Several TREM2-activating antibodies, including AL002 (Alector/AbbVie), 4D9 (Denali Therapeutics), and VGL101 (Vigil Neuroscience), have progressed into clinical development (Schlepckow, Monroe et al. 2020, Wang, Mustafa et al. 2020). These antibodies demonstrate promising efficacy in preclinical models by enhancing microglial activation and facilitating clearance of pathological aggregates. However, antibody-based therapeutics face inherent limitations including high manufacturing costs, potential immunogenicity, limited blood-brain barrier (BBB) penetration, and complex dosing regimens requiring intravenous administration (Pardridge 2012). In contrast, small-molecule TREM2 agonists offer distinct advantages including oral bioavailability, superior BBB permeability, and cost-effective manufacturing. VG-3927, developed by Vigil Neuroscience, represents the first small-molecule TREM2 agonist to enter clinical trials, demonstrating high selectivity for membrane-bound TREM2 and favorable brain penetration (Mirescu 2024). Despite this progress, current small-molecule approaches face significant challenges related to selectivity and comprehensive target modulation. A critical limitation of existing TREM2 agonists is their inability to address ADAM (A Disintegrin And Metalloproteinase)-mediated receptor shedding. TREM2 undergoes constitutive proteolytic cleavage by ADAM10 and ADAM17 at the His157-Ser158 peptide bond, generating soluble TREM2 (sTREM2) and reducing functional receptor levels on the cell surface (Kleinberger, Yamanishi et al. 2014, Heslegrave, Heywood et al. 2016, Feuerbach, Schindler et al. 2017). This shedding process significantly diminishes TREM2 signaling capacity and therapeutic efficacy (Suárez-Calvet, Kleinberger et al. 2016). While several antibodies, including 4D9, have demonstrated anti-shedding properties by sterically protecting the cleavage site, no small-molecule agonist has been reported to possess this dual functionality (Heslegrave, Heywood et al. 2016, van Lengerich, Zhan et al. 2023). Additionally, selectivity between TREM family members remains a concern. TREM1, another member of the TREM family, is primarily expressed on neutrophils and monocytes and functions as an amplifier of inflammatory responses (Bouchon, Facchetti et al. 2001). Cross-reactivity with TREM1 could potentially lead to unwanted pro-inflammatory effects, particularly in the context of chronic neuroinflammation (Derive, Massin et al. 2010, Singh, Rai et al. 2021).

To overcome these therapeutic limitations, we employed Affinity Selection-Mass Spectrometry (AS-MS) as an unbiased discovery platform for identifying novel TREM2 modulators. Unlike conventional functional screens that may overlook compounds with non-traditional mechanisms, AS-MS enables direct detection of target engagement, facilitating discovery of molecules with unprecedented modes of action (Dueñas, Peltier-Heap et al. 2023). This approach has been successfully employed in other challenging targets, including STING (Stimulator of Interferon Genes), where Affinity Selection-Mass Spectrometry (AS-MS) screening identified novel agonists from large chemical libraries (Decout, Katz et al. 2021, Muchiri and van Breemen 2021). The integration of MS-based primary screening with orthogonal biophysical validation methods, including Temperature-Related Intensity Change (TRIC) analysis via Dianthus platforms, Microscale Thermophoresis (MST), and Surface Plasmon Resonance (SPR), provides a comprehensive framework for hit identification and characterization (Zhang, Calvo-Barreiro et al. 2024, Zhao, Hadavi et al. 2024). This multi-tiered approach ensures both binding specificity and kinetic validation, while enabling detailed characterization of binding affinity, protein stability, and mechanistic insights.

Through the implementation of Affinity Selection-Mass Spectrometry (AS-MS) as our primary discovery platform, we identified As48, a novel small-molecule compound that exhibits unprecedented dual functionality as both a TREM2 agonist and ADAM shedding inhibitor. AS-MS enabled unbiased detection of direct TREM2 binders from chemical libraries, providing a critical advantage over conventional functional screens that might miss compounds with novel mechanisms of action. Beginning with AS-MS screening of small molecules, we identified 62 potential TREM2 binders, which were subsequently validated through secondary screening using TRIC-based binding analysis. Lead compounds were further characterized using MST and SPR to determine binding kinetics and confirm target engagement. Comprehensive cellular and biochemical characterization revealed that As48 possesses several unique properties that distinguish it from existing TREM2 modulators. Notably, As48 demonstrates high selectivity for TREM2 over TREM1, potentially avoiding unwanted pro-inflammatory side effects. Most significantly, As48 represents the first small-molecule TREM2 agonist capable of simultaneously inhibiting ADAM-mediated receptor shedding, thereby preserving surface receptor levels while enhancing downstream signaling. This dual mechanism of action positions As48 as a potentially superior therapeutic approach for neurodegenerative diseases characterized by TREM2 dysfunction. By combining receptor activation with protection from proteolytic degradation, As48 may achieve more robust and sustained therapeutic effects compared to traditional agonist-only approaches. The identification of this novel dual-function modulator not only advances our understanding of TREM2 pharmacology but also establishes a new paradigm for developing next-generation neuroinflammatory therapeutics.

## Methods

### Cell culture

HMC3 human microglial cells were obtained from ATCC (Manassas, VA, USA) and cultured in Eagle’s Minimum Essential Medium (EMEM) supplemented with 10% fetal bovine serum (FBS), 100 U/mL penicillin, 100 μg/mL streptomycin, 2 mM L-glutamine, and 1 mM sodium pyruvate. Cells were maintained at 37°C in a humidified atmosphere containing 5% CO₂. For the generation of TREM2-overexpressing HMC3 cells, a lentiviral transduction system was employed using the pLenti6P.3-hTREM2 vector. Stable cell lines were selected and maintained using 1 μg/mL puromycin.

### Protein purification

The human TREM2 ectodomain was cloned into a pCMV3 vector with an N-terminal His₆ tag and a 3C protease cleavage site. The construct was transiently transfected into Expi293F™ cells using the ExpiFectamine™ 293 Transfection Kit, following the manufacturer’s protocol. Cells were cultured in Expi293™ Expression Medium at 2.5 × 10⁶ cells/mL in vented flasks and incubated at 37°C, 8% CO₂, 100 rpm for 5 days (Szykowska, Chen et al. 2021). The culture medium was clarified by centrifugation at 4,000 × g for 15 minutes at 4°C and filtered through a 0.22 µm membrane. The supernatant was incubated with Ni-NTA agarose beads (Qiagen) pre-equilibrated with binding buffer (20 mM HEPES pH 7.5, 150 mM NaCl, 20 mM imidazole and 5% Glycerol). After washing with 10 column volumes of the same buffer, the bound protein was eluted with 300 mM imidazole in the same buffer base (Upadhyay, Bhardwaj et al. 2025). Eluted fractions were pooled and further purified by size-exclusion chromatography using a Superdex 200 Increase 10/300 GL column (Cytiva) equilibrated in 20 mM HEPES pH 7.5, 150 mM NaCl, 5% Glycerol (Khan, Upadhyay et al. 2024). Monomeric fractions were collected, concentrated to 5–10 mg/mL, and used directly for binding assays.

### Affinity Selection-Mass Spectrometry (AS-MS) Screening

AS-MS screening was performed as previously described (Zhang, Calvo-Barreiro et al. 2024). Briefly, recombinant human TREM2 protein was immobilized on magnetic beads and incubated with small molecule libraries. After washing to remove non-specific binders, bound compounds were eluted and analyzed by liquid chromatography-mass spectrometry (LC-MS). Compounds showing specific binding to TREM2 versus control beads were selected for secondary validation. The AS-MS approach enabled identification of direct TREM2 binders without relying on functional readouts, providing an unbiased discovery platform for novel mechanisms.

### Temperature-Related Intensity Change (TRIC) Binding Assay

Temperature-Related Intensity Change (TRIC) measurements were performed using a Dianthus NT.23PicoDuo instrument (NanoTemper Technologies, Munich, Germany). Recombinant human TREM2 protein (in-house) was labeled with RED-tris-NTA 2nd Generation dye using the His-Tag Labeling Kit (NanoTemper Technologies) according to the manufacturer’s protocol. All binding experiments were conducted in PBST buffer containing 154 mM NaCl, 5.6 mM Na₂HPO₄, 1.05 mM KH₂PO₄, pH 7.4, and 0.005% Tween-20. For binding measurements, labeled TREM2 protein was used at a final concentration of 10 nM. Test compounds were serially diluted in PBST buffer and incubated with labeled TREM2 for 10 minutes at room temperature (22-25°C) prior to analysis. The Dianthus instrument was configured with the following parameters: 85% LED excitation power, picomolar detector sensitivity disabled, and laser on-time of 5 seconds. Data analysis was performed using Dianthus Analysis software (NanoTemper Technologies) for initial processing, followed by curve fitting and statistical analysis using GraphPad Prism 10.0 (GraphPad Software, San Diego, CA, USA).

### Microscale Thermophoresis (MST) Binding Assay

Binding affinity measurements were performed using microscale thermophoresis (MST) on a Monolith NT.115 system (NanoTemper Technologies, Munich, Germany). Recombinant human TREM2 protein (in-house) and human TREM1 protein (BioTechne, Minneapolis, MN, USA) were labeled using the RED-tris-NTA His-tag labeling kit (NanoTemper Technologies) according to the manufacturer’s instructions. MST measurements were conducted using different buffer conditions optimized for each protein: TREM2 assays were performed in PBS buffer (pH 7.4) containing 0.005% Tween-20, while TREM1 assays used PBS buffer (pH 7.4) with 0.05% Tween-20. Labeled proteins were used at a final concentration of 40 nM and incubated with serially diluted test compounds for 10 minutes at room temperature (22-25°C) prior to measurement. MST experiments were performed using standard capillaries with the following instrument parameters: red filter set, 100% LED power, and medium MST power. Thermophoresis was monitored for 20 seconds with an additional 5-second delay. Data analysis was conducted using MO.Affinity Analysis software (NanoTemper Technologies) for initial processing, followed by curve fitting using GraphPad Prism 10.0 (GraphPad Software, San Diego, CA, USA).

### Thermal Shift Assay (TSA)

Thermal stability of recombinant human TREM2 protein was assessed using the Prometheus Panta system (NanoTemper Technologies, Munich, Germany). TREM2 protein (100 nM) was incubated with As48 (30 μM) or DMSO control (1%) in PBS buffer (pH 7.4) for 20 min at room temperature. Samples were loaded into Prometheus^TM^ NT.48 capillaries and subjected to a temperature gradient from 40°C to 95°C at a heating rate of 1°C/min. Melting temperatures (Tm) were determined by monitoring intrinsic tryptophan fluorescence changes (330 nm and 350 nm). Data were analyzed using Panta Control software.

### Surface plasmon resonance (SPR)

The interaction between As48 and the human TREM2 protein was evaluated using surface plasmon resonance (SPR) on a Biacore 8K instrument (Cytiva). Biotinylated TREM2 (Cat. No. 11084-H49H-B, Sino Biological) was immobilized on an SA Sensor Chip (Cytiva) in PBS-P buffer (0.2 M phosphate buffer, 27 mM KCl, 1.37 M NaCl, and 0.5% Surfactant P20, pH 7.4; Cytiva) to a response level of RU 4,343 ± 399 RU. Before protein immobilization, the chip surface was conditioned with three 1-minute injections of 1 M NaCl in 50 mM NaOH. Following immobilization, the surface was washed sequentially with 50% (v/v) isopropanol in water, 1 M NaCl, and 50 mM NaOH to remove weakly or non-specifically bound material. The system was then equilibrated in running buffer until a stable baseline was established. Kinetic measurements were performed using a single-cycle kinetic method. Five concentrations of As48 (100, 33.33, 11.11, 3.70, and 1.23 μM) were prepared in PBS-P buffer containing 2% DMSO (Cytiva) and injected over the immobilized TREM2 surface. Experiments were carried out at 25°C with a flow rate of 30 μL/min, a contact time of 120 s, and a dissociation phase of 600 s. After each run, the surface was washed with 50% DMSO. Sensorgram was generated after subtracting background response signal from a reference flow cell and from a control experiment with buffer injection. Interaction was investigated at least in triplicate. Data analysis was performed using Biacore™ Insight Evaluation Software (Cytiva). A 1:1 binding model provided the best fit to the experimental data in single-cycle kinetic analysis.

### Cell viability assay

The cytotoxic effects of test compounds were evaluated using the MTS assay (Cell Titer 96® AQueous One Solution, Promega, WI, USA). HMC3 microglial cells were seeded in 96-well plates and maintained in complete growth medium containing 10% FBS until reaching ap-proximately 40% confluence (24 h). Cells were then treated with compounds and Tween-20 for overnight followed by MTS analysis according to the supplier’s protocol. Absorbance readings were obtained at 490 nm using an Infinite M1000 Pro Microplate Reader (Tecan, Männedorf, Switzerland).

### AlphaLISA

The Phospho-AlphaLISA assay (Revvity, USA) was employed to quantify the phosphorylation of the spleen tyrosine kinase (Syk) at a specific residue. (Y525/526) This bead-based immunoassay utilizes two antibodies: one specific for the phosphorylated epitope and the other recognizing a distal epitope on the same protein. Upon binding to the phosphorylated target, proximity of donor and acceptor beads induces a luminescent Alpha signal. The signal intensity is directly proportional to the concentration of phosphorylated protein in the sample. HEK-hTREM2/DAP12 cells were seeded at a density of 5 × 10⁴ cells per well in 96-well plates (100 µL/well) in Dulbecco’s Modified Eagle Medium (DMEM; Gibco) supplemented with 10% fetal bovine serum (FBS; Gibco). Cells were incubated for 24 hours at 37 °C in a humidified atmosphere containing 5% CO₂. Compounds VG-3927 and AS48 were prepared in complete media at a final concentration of 5, 10, 25, 50, 75 or 100 µM. Culture medium was aspirated from each well and replaced with compound-containing medium. Vehicle control wells received the solvent used for compound dilution (DMSO). Following specific times of incubation at 37 °C, the medium was gently removed, and cells were lysed using the lysis buffer provided in the assay kit. After complete lysis, the Phospho-AlphaLISA assay was performed according to the manufacturer’s instructions.

### Intracellular Calcium Measurement

HEK293 cells were grown on 96-well clear bottom black wall plates (Corning Inc., Corning, NY, USA). The intracellular calcium was measured using the Fluo-4 NW kit (Invitrogen, Carlsbad, CA, USA) following the manufacturer’s instructions. Briefly, cells were incubated with 100 µL assay buffer with Fluo 4 dye for 1 h. The fluorescence was measured using the EVOS® FL Auto Imaging System (Thermo Scientific, Waltham, MA, USA)

### Immunoblotting

For Western blot analysis, HEK293 and HMC3 cell were plated on 6-well-plates and incubated overnight. Cells were treated with compounds accordingly, washed twice with ice-cold PBS, and lysed for 15 min in RIPA buffer supplemented with a protease inhibitor cocktail. Lysed samples were centrifuged at 13,000 rpm for 20 min at 4 °C. Extracted proteins were quantified using the Bradford protein assay kit (Thermo Scientific, Waltham, MA, USA), and 40 µg of total proteins was loaded to each well and separated by 4–20% Tris-glycine precast gel (Bio-rad, PA, USA). Proteins were transferred to PVDF membranes (Bio-rad, PA, USA), followed by blocking for 1 h with SuperBlock™ Blocking Buffer (Thermo Scientific, Waltham, MA, USA). The membranes were incubated with primary antibodies overnight at 4 °C with the indicated primary antibodies; anti-pSYK (Cell Signaling Technology, MA, USA, Cat#2711), anti TREM2 (extracellular domain) (Thermo Scientific, Waltham, MA, USA, Cat#4H42L9), anti TREM2 (Cell Signaling Technology, MA, USA, Cat#D8I4C) and anti-β-actin (Cell Signaling Technology, MA, USA, Cat#13E5). Then, the membranes were washed three times in TBST and incubated with horseradish peroxidase-conjugated secondary antibodies for 1 h. After being washed three times, membranes were detected using the SuperSignal^TM^ West Pcio PLUS detection system (Thermo Scientific, Waltham, MA, USA). All experiments were repeated 3 times independently, and ImageJ software (NIH, Bethesda, MD, USA) was used for result analysis.

### Phagocytosis Assay in BV2 Cells

Phagocytic activity in BV2 microglial cells was assessed using green fluorescent latex beads (Sigma, USA). BV2 cells were serum-starved, then treated with either compound VG-3927 or AS48 (25 µM) for 30 minutes, or with the corresponding solubilization buffer (DMSO) as a control. Following treatment, beads were added to the culture medium and incubated for an additional 30 minutes at 37 °C. Cells were washed with ice-cold phosphate-buffered saline (PBS) prior fixation with 4% paraformaldehyde (PFA; ThermoFisher, USA) and processed for immunocytochemistry. Images were acquired under fluorescent microscope and analyzed. Cells positive for phagocytosis were defined as IBA1-positive cells containing at least one internalized fluorescent bead.

### Immunocytochemistry

Fixed cells were permeabilized with Triton X-100-containing PBS and subsequently blocked with 1% BSA. Immunostaining was then performed using primary anti IBA1 (Wako, Richmond, VA) antibody followed by an Alexa Fluor 594-conjugated secondary antibody (ThermoFisher).

### Molecular docking simulation

To investigate the binding mode and dynamic stability of As48 with TREM2, a combined molecular docking and molecular dynamics simulation approach was employed. The three-dimensional structure of human TREM2 (PDB ID: 6Y6C)(Szykowska, Chen et al. 2021) was prepared using the Protein Preparation Wizard in Maestro (Schrödinger Suite V.2023.2.). The protein preparation step involved addition of hydrogens, optimization of hydrogen-bonding networks, and energy minimization using the OPLS4 force field. Given the reported conformational flexibility of TREM2, the Induced Fit Docking protocol in Maestro was chosen for carrying out the docking assay to allow side-chain and backbone adjustments upon ligand binding (Dean, Roberson et al. 2019, Mai, Wei et al. 2022). As48 was prepared using the LigPrep module within Maestro to generate low-energy conformers with appropriate protonation states at physiological pH (Nada, Zhang et al. 2025). The binding site was defined around the His-tag interface, where cleavage has been previously reported and docking was carried out with default precision settings (Schlepckow, Monroe et al. 2020). The top-ranked complex based on Glide docking scores and Prime energy refinements was selected for further analysis. Discovery studio Visualizer V2024.1. was employed for the Visualization of the docking results.

To evaluate the stability of the docked complex and investigate ligand-induced conformational effects, MD simulations were carried out using Desmond (Schrödinger). The protein–ligand complex was embedded in an orthorhombic TIP3P water box with a 10 Å buffer, and appropriate counterions were added to neutralize the system. The system was subjected to energy minimization followed by equilibration using the default relaxation protocol. A 100 ns production run was performed in the NPT ensemble at 300 K and 1 atm, with periodic boundary conditions, using the OPLS4 force field and RESPA integrator. Another independent simulation was carried out for the unbound TREM2 protein. The simulation trajectory was analyzed to compute the root-mean-square deviation (RMSD) and root-mean-square fluctuation (RMSF) of the protein backbone and Cα atoms. Protein–ligand interactions, including hydrogen bonding, hydrophobic contacts, and water bridges were evaluated over the 100ns using the Simulation Interaction Diagram tool in Maestro to identify key residues contributing to ligand stability and potential inhibition of TREM2 cleavage. Xmgrace was used to plot the RMSD results for the unbound TREM2 and its complex with As48.

### ADAM10 and ADAM17 Proteolytic Activity Measurements

ADAM10 and ADAM17 proteolytic activities were measured using ADAM10 Fluorogenic Assay Kit and ADAM17 Fluorogenic Assay Kit, respectively (BPS Bioscience, San Diego, CA, USA). All assays were performed according to the manufacturer’s protocol. For activity inhibition studies, recombinant ADAM10 or ADAM17 enzymes were pre-incubated with test compounds at indicated concentrations for 30 minutes at room temperature prior to substrate addition. GM6001, a broad-spectrum metalloproteinase inhibitor, was used as a positive control for enzyme inhibition. As48 was tested to evaluate its potential effects on ADAM protease activity. Fluorogenic substrate cleavage was monitored using an Infinite M1000 Pro Microplate Reader (Tecan, Männedorf, Switzerland) with excitation at 485 nm and emission at 530 nm. All experiments were performed in triplicate.

### Pharmacokinetics (PK) study

The preliminary evaluation of PK parameters for As48 was performed as previously reported by us (Kaur, Nada et al. 2025, Nada, Zhang et al. 2025). In brief, physicochemical and biological profiling included determination of LogD₇.₄, microsomal stability, kinetic solubility, and cytotoxicity across a panel of cell lines. Solubility was assessed using UV–vis spectrophotometry, while cell viability was measured using the PrestoBlue® assay (ThermoFisher, Cat# A13261).

## Results

### Expression and Purification of Recombinant Human TREM2 Protein

Recombinant hTREM2 (19–174) was efficiently expressed in Expi293F™ cells and purified using a two-step procedure. The Ni-NTA purification yielded protein of high purity, which was confirmed by a single broad band on SDS-PAGE under reducing conditions, migrating between 30–40 kDa—higher than the theoretical mass of 19.3 kDa due to expected glycosylation. Subsequent size-exclusion chromatography on a Superdex 200 Increase 10/300 GL column revealed a single, symmetric peak eluting at ∼15 mL. This elution volume corresponds to a monomeric species, with no evidence of aggregates or higher-order oligomers. The purified protein was stable in HEPES–NaCl–glycerol buffer and was directly used for downstream binding experiments. The SEC profile is shown in (Fig 1B).

**Figure 1.**
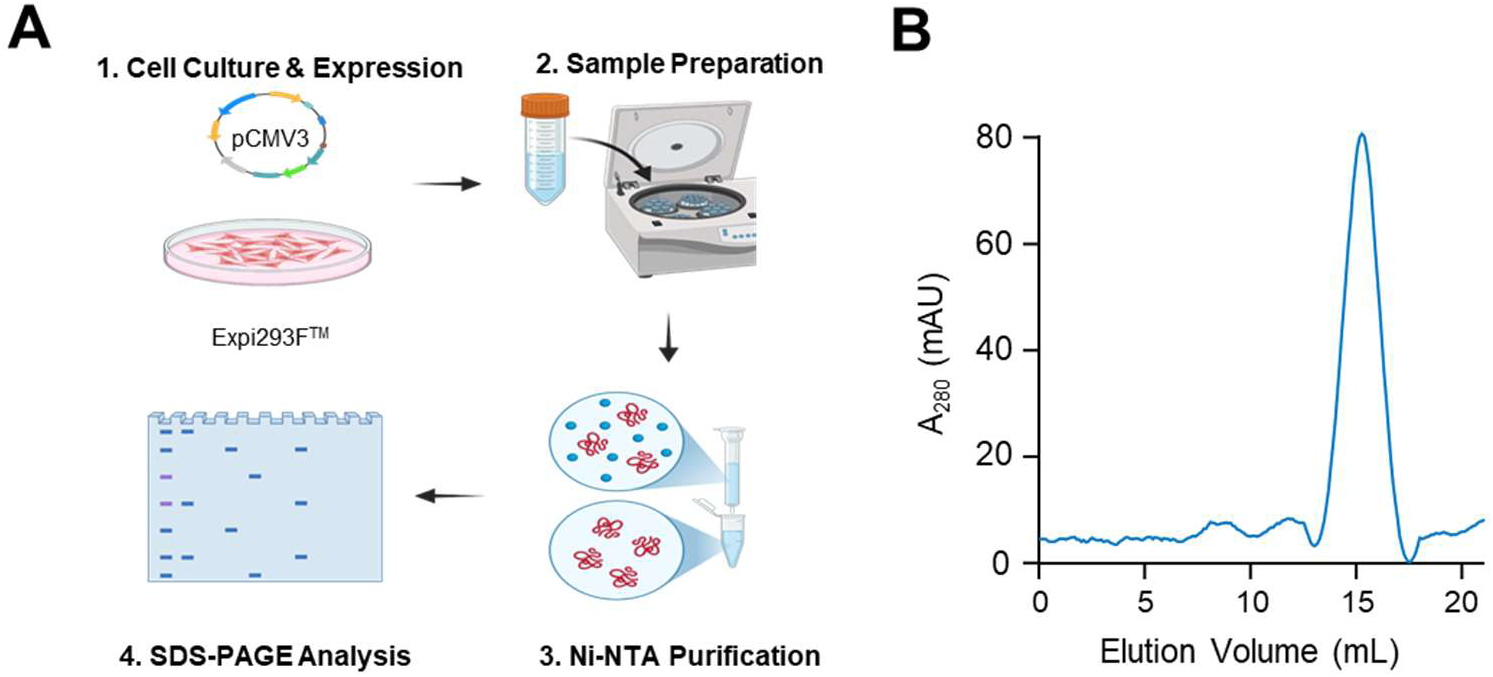
Expression and purification workflow of recombinant human TREM2 protein. **A)** Schematic workflow for human TREM2 ectodomain (residues 19-174) production. **B)** Size-exclusion chromatography profile of purified TREM2.

### Mass spectrometry guided discovery and validation of TREM2 binding compounds

To identify novel TREM2-binding compounds, we employed Affinity Selection-Mass Spectrometry (AS-MS) as our primary discovery platform. This unbiased approach enabled direct detection of TREM2 binders from chemical libraries without relying on functional readouts. AS-MS screening identified 62 potential TREM2 binders, which were subsequently validated using secondary screening platforms. Single-dose screening was performed using the Dianthus platform, which utilizes Temperature-Related Intensity Change (TRIC) technology to detect direct protein-compound interactions. For this screening, compounds were tested at 100 μM concentration against recombinant human TREM2 protein (10 nM). PC-192, a known TREM2 agonist, served as the positive control to validate assay performance. We established a stringent hit selection criterion, defining TREM2-binding compounds as those exhibiting signal changes greater than 5-fold the standard deviation of the negative control (TREM2 alone). From the 63 compounds screened, we successfully identified two promising TREM2-binding hits: As37 and As48 (Fig 2A). The remaining compounds either showed no significant binding activity or produced false-positive signals attributed to compound aggregation or fluorescence interference. To determine the relative binding affinities and validate these hits, we conducted concentration-dependent TREM2 binding experiments using TRIC analysis. As37 demonstrated moderate binding affinity with an equilibrium dissociation constant (K_D_) of 95.53 ± 7.94 μM (Fig 2B). In contrast, As48 exhibited significantly stronger binding affinity with a K_D_ of 12.48 ± 2.48 μM (Fig 2C). Based on these comparative binding studies, As48 was selected as our primary lead compound for further biophysical characterization and functional validation due to its superior binding affinity to TREM2 protein.

**Figure 2.**
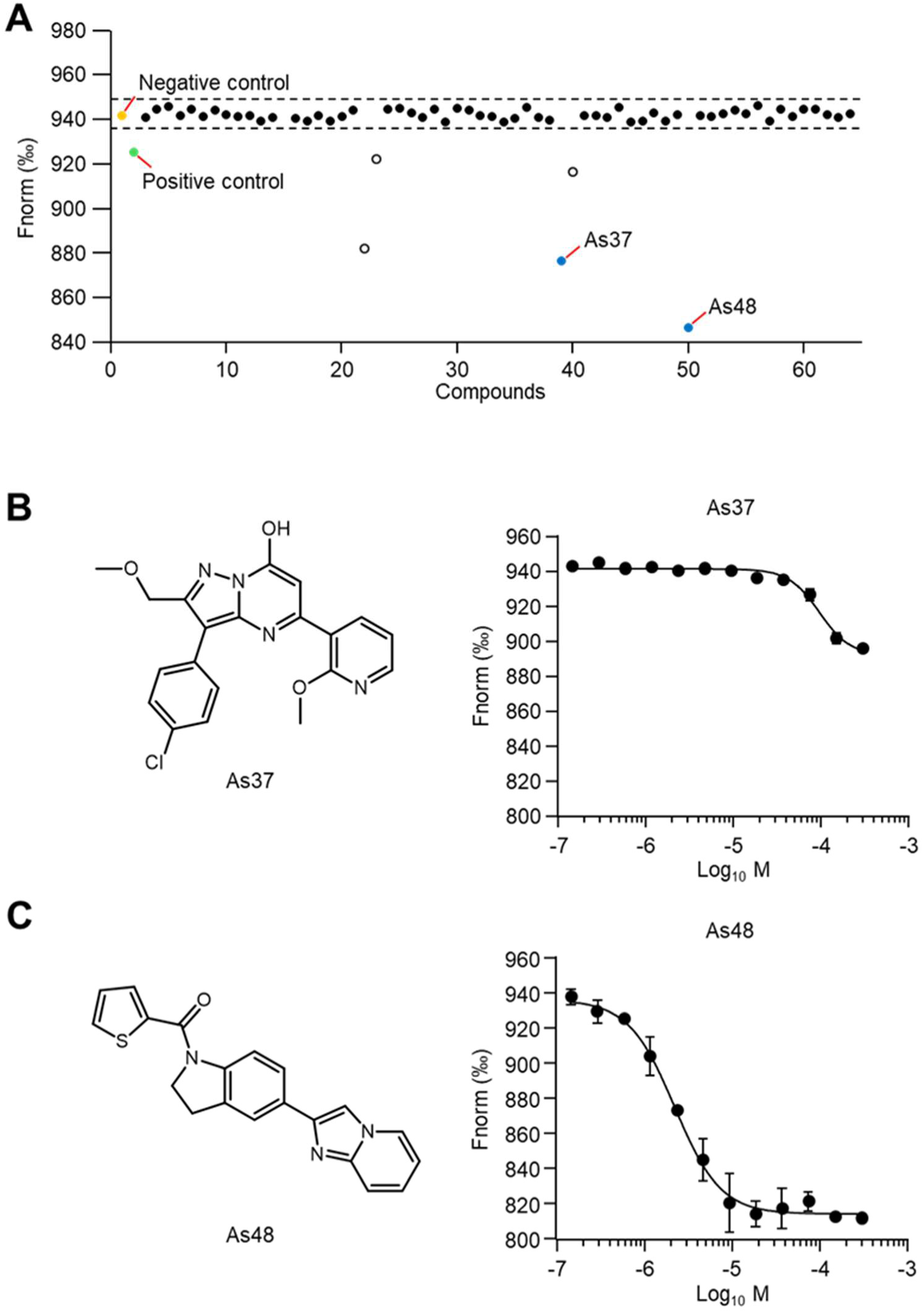
Identification and validation of TREM2-binding compounds through MS-guided screening and biophysical characterization. **A)** Single-concentration screening of 63 compounds against TREM2 protein (10 nM) at 100 μM using Temperature-Related Intensity Change (TRIC) analysis. Yellow dot: negative control (TREM2 alone); green dot: positive control (TREM2 with PC-192). The dashed line represents the threshold for hit identification, corresponding to signals exceeding 5-fold the standard deviation of the negative control. Blue dots indicate validated TREM2-binding hits (As37 and As48), while white dots represent false positives due to compound aggregation or interference. **B)** Chemical structure of As37 (left panel) and its concentration-dependent binding profile determined by TRIC analysis (right panel). **C)** Chemical structure of As48 (left panel) and its concentration-dependent binding assessment by TRIC analysis (right panel). Data are presented as mean ± SEM (n = 3).

### Comprehensive biophysical validation of As48-TREM2 binding

To establish definitive evidence for direct As48-TREM2 interaction, we used multiple biophysical techniques that collectively confirm binding through different detection principles. Microscale thermophoresis (MST) measurements provided quantitative binding affinity data, revealing that As48 binds to TREM2 with a KD of 13.8 ± 1.08 μM (Fig 3A). This value is consistent with our initial TRIC-based screening results, confirming the reliability of our binding measurements across different platforms.

**Figure 3.**
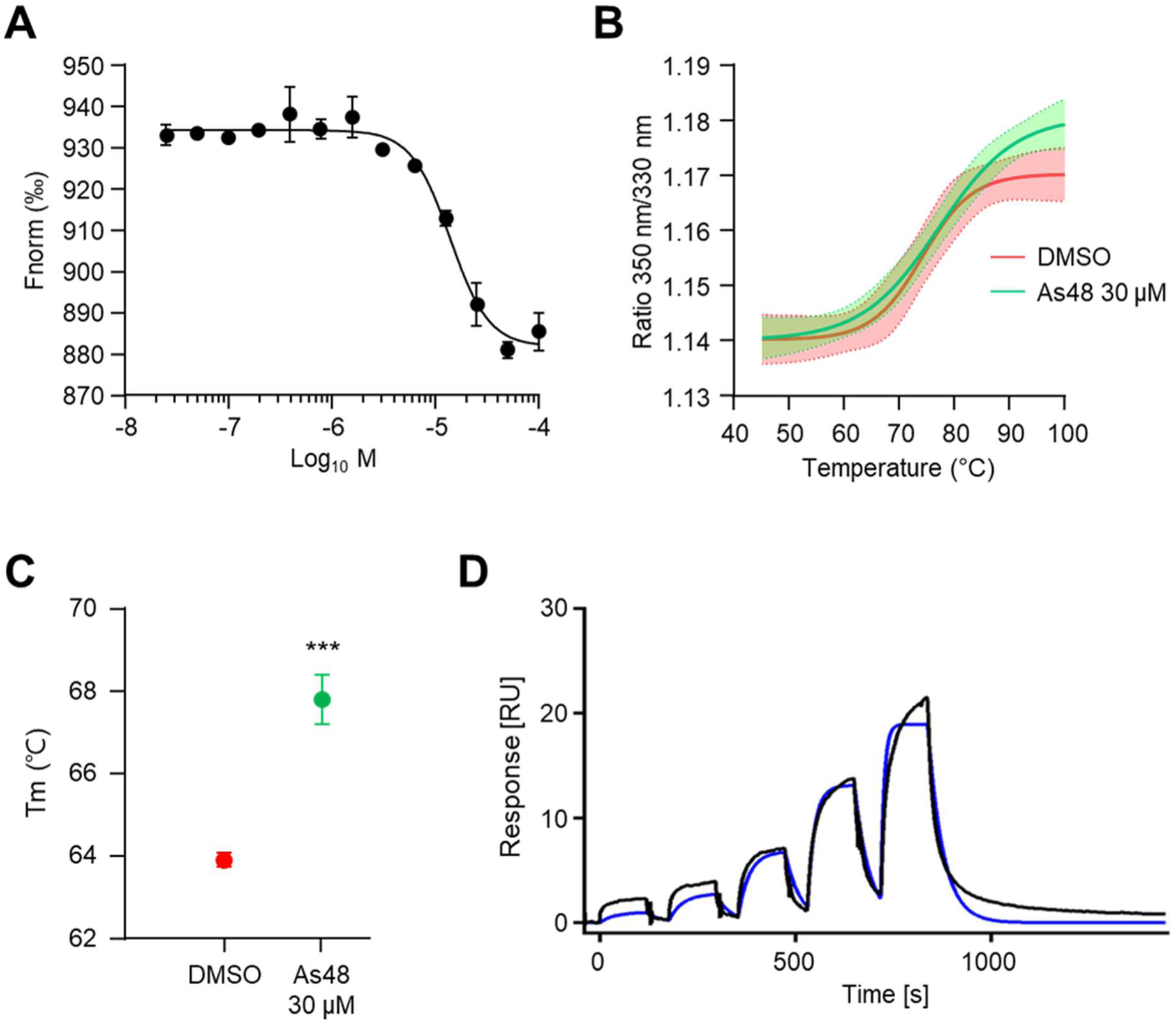
Comprehensive biophysical characterization of As48 TREM2 binding interactions. **A)** Dose-dependent binding experiments for As48 using MST. **B)** Thermal stability analysis of TREM2 protein using Prometheus Panta system. Raw melting curves showing the ratio of fluorescence signals (350 nm/330 nm) as a function of temperature (45-100°C) for DMSO control (red) and As48-treated samples (30 μM, green). Shaded areas represent SEM. **C)** Melting temperature (Tm) comparison between DMSO control and As48 treatment (30 μM). ***P < 0.001 analyzed by Student’s unpaired two-tailed t test. **D)** Single-cycle kinetic sensorgrams of As48 interacting with the TREM2 protein using SPR. Black lines represent the experimental data, while blue lines show the 1:1 kinetic binding model fit.

To evaluate whether As48 binding induces structural stability in TREM2, we performed thermal shift assays using the Prometheus Panta system. This technique monitors protein thermal stability by tracking intrinsic tryptophan fluorescence changes during controlled heating (Niesen, Berglund et al. 2007). As48 treatment (30 μM) significantly increased the melting temperature of TREM2 from 63.93 ± 0.14 ℃ (DMSO control) to 67.79 ± 0.61 ℃, representing a substantial thermal shift of +3.86 ℃ (Fig 3B and C). This pronounced stabilization strongly indicates that As48 binding induces conformational changes that enhance protein thermal stability, providing compelling evidence for direct protein-compound interaction.

To obtain comprehensive binding kinetics data, we employed surface plasmon resonance (SPR) analysis for real-time monitoring of As48-TREM2 interactions. Recombinant TREM2 protein was immobilized on the sensor chip surface, and serial dilutions of As48 were injected to generate concentration-dependent binding profiles. The resulting sensorgrams demonstrated clear association and dissociation phases, confirming specific binding interactions (Fig 3D). Kinetic analysis using a 1:1 binding model yielded a K_D_ of 29.5 μM, which, while slightly higher than MST measurements, confirms the micromolar binding affinity range and validates the direct interaction between As48 and TREM2. The convergent results from these three independent biophysical techniques provide robust evidence for specific, direct binding of As48 to TREM2 protein, with binding affinities consistently in the low-to-mid micromolar range across all platforms.

### Selectivity and cytotoxicity assessment of As48

To evaluate the selectivity profile of As48, we assessed its binding affinity to TREM1, a closely related family member that shares structural similarities with TREM2 but has distinct biological functions. MST analysis revealed that As48 binds to TREM1 with a K_D_ of 97.92 ± 23.12 μM (Fig 4A), which represents approximately 7-fold weaker binding compared to its affinity for TREM2 (K_D_ = 13.8 ± 1.08 μM). Given that TREM2 is predominantly expressed in microglial cells, we evaluated the cytotoxicity profile of As48 using HMC3 human microglial cells, a well-established cell line model for studying microglial biology. Cells were treated with increasing concentrations of As48 (0-300 μM) for overnight, and cell viability was assessed using the MTS colorimetric assay. Remarkably, As48 demonstrated showing no significant cytotoxicity even at concentrations up to 100 μM (Fig 4B). Even at the highest tested concentration of 300 μM, more than 20-fold above its TREM2 binding affinity, As48 induced only modest cytotoxicity (∼20% reduction in cell viability). These results establish that As48 has a favorable pharmacological profile characterized by selective binding to TREM2 over TREM1 and minimal cytotoxicity in the target cell population, supporting its potential as a safe and selective TREM2 modulator for further development.

**Figure 4.**
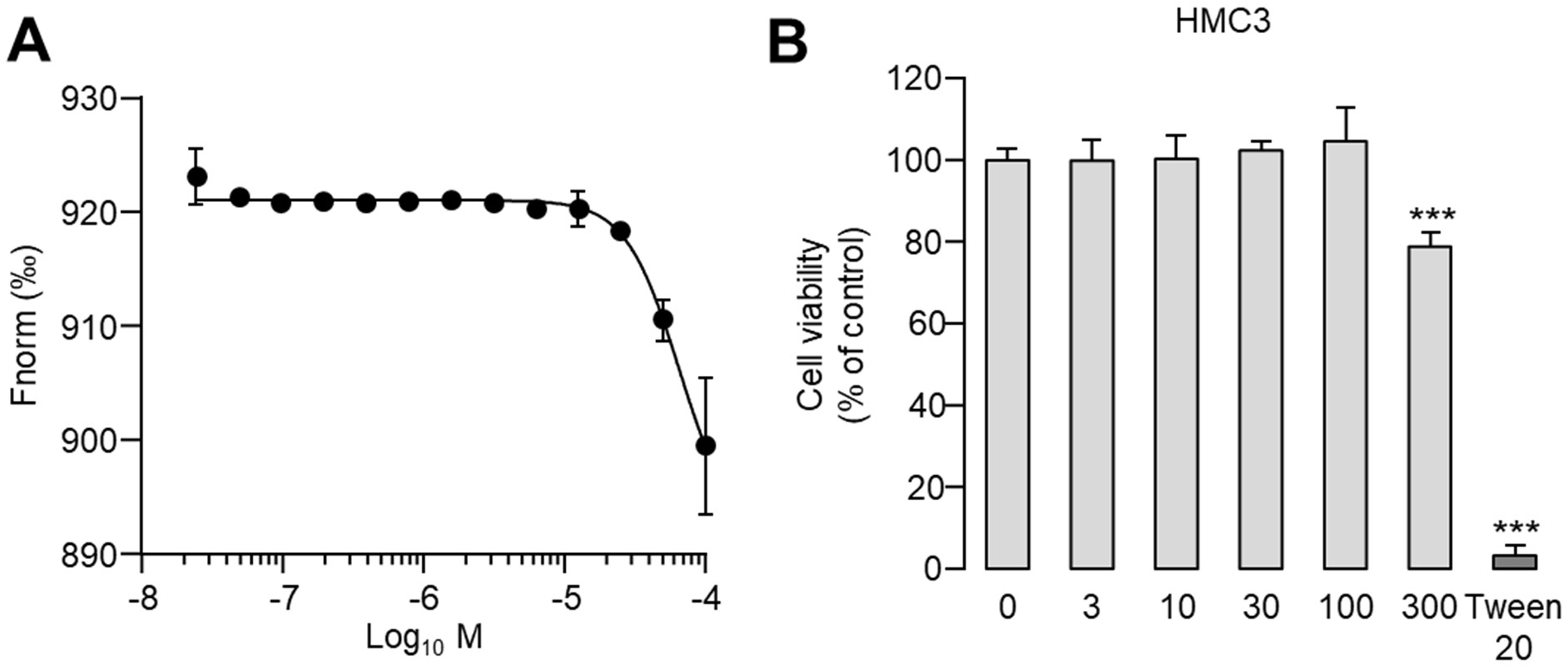
Assessment of As48 selectivity and cytotoxicity profile. **A)** Dose-dependent binding experiments for As48 in TREM1 using MST. Mean ± SEM (n = 3). **B)** Cytotoxicity assessment of As48 in HMC3 human microglial cells. Cells were treated with indicated concentrations of As48 (0-300 μM) for over nights, and cell viability was determined using MTS colorimetric assay. Tween 20 (0.1%) served as a positive control for cytotoxicity. Data are presented as mean ± SEM (n = 5). Statistical significance was determined by unpaired two-tailed t-test (***p < 0.001).

### As48 functions as a TREM2 agonist by inducing SYK phosphorylation and downstream calcium signaling

To assess the agonistic activity of As48 on TREM2 signaling, we employed HEK293 cells stably expressing human TREM2 and its signaling adaptor DAP12 (HEK293-hTREM2/DAP12), using SYK phosphorylation as a proximal readout of TREM2 activation. In a time-course analysis, VG-3927 (25 μM) induced rapid SYK phosphorylation detectable at 5 minutes, with levels continuing to increase and peaking at 60 minutes. In contrast, As48 (25 μM) triggered a delayed yet sustained response, initiating at 15 minutes and remaining elevated through 60 minutes (Fig 5A). Dose-response analysis revealed that VG-3927 elicited SYK activation starting at 5 μM, with maximal phosphorylation observed at 25 μM, while higher concentrations led to reduced activity. As48 displayed a higher activation threshold, with SYK phosphorylation detected from 10 μM and maximal activity at 25 μM (Fig 5B).

**Figure 5.**
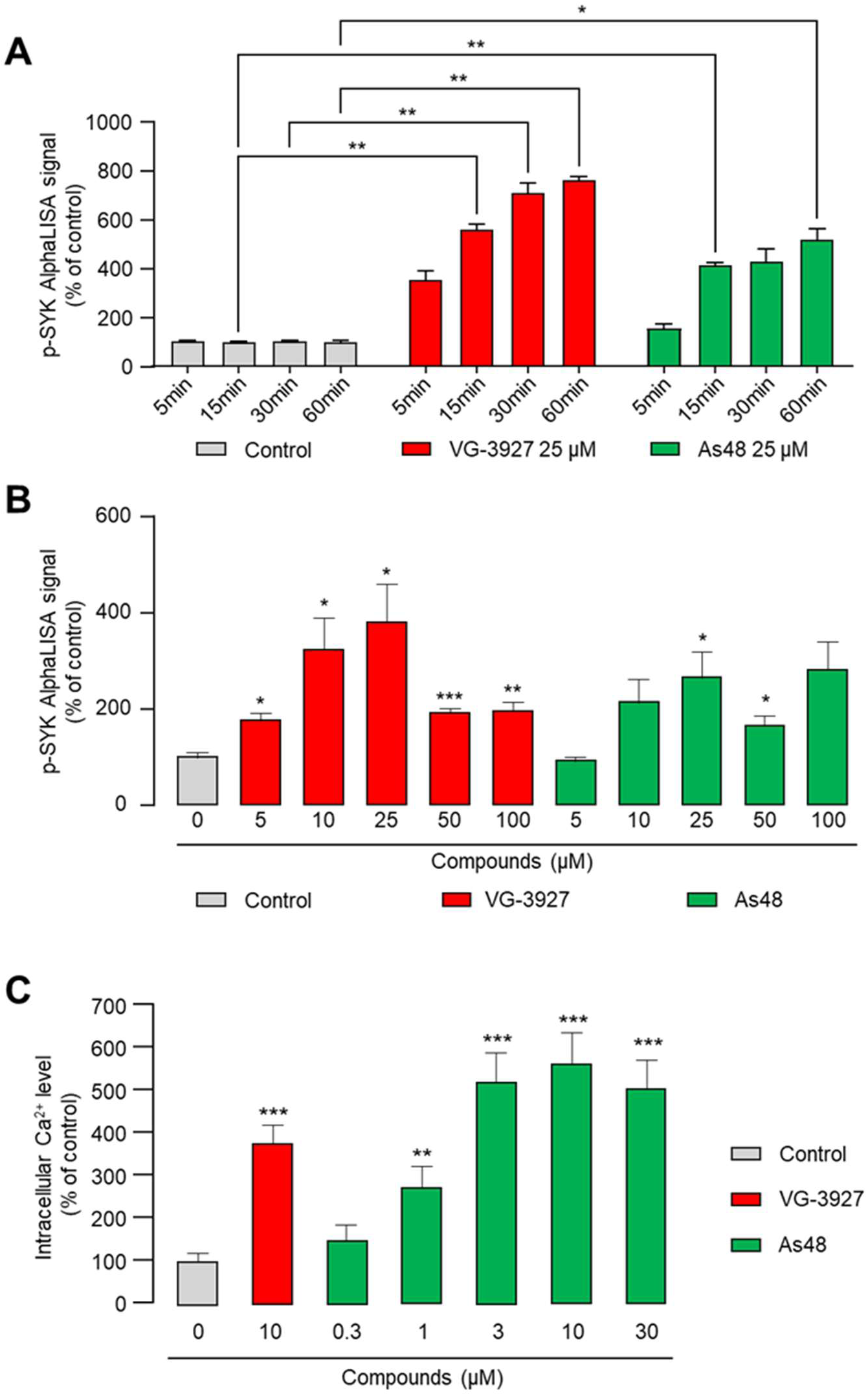
As48 activates TREM2 signaling through SYK phosphorylation and calcium signaling in HEK293-hTREM2/DAP12 cells. **A)** Time-course analysis of SYK phosphorylation in HEK293-hTREM2/DAP12 cells treated with compounds. Cells were stimulated with vehicle control (DMSO, gray bars), VG-3927 (25 μM, red bars), or As48 (25 μM, green bars) for 5, 15, 30, or 60 minutes. Phosphorylated SYK levels were quantified using an AlphaLISA assay and normalized to vehicle-treated controls. Data represent mean ± SEM from 4 independent experiments. Statistical significance was assessed using two-way Anova followed by Tukey’s multiple comparison test (*p<0.05, **p < 0.01, ***p < 0.001). **B)** Dose-dependent induction of SYK phosphorylation. Cells were treated for 60 minutes with increasing concentrations of VG-3927 (red bars) or As48 (green bars), and phospho-SYK levels were quantified as in **A)**. DMSO-treated cells (gray bar) served as the baseline control. Data represent mean ± SEM from 3 independent experiments (n=6 per conditions). Statistical significance was assessed using mixed effects model analysis followed by Tukey’s multiple comparison test (**p < 0.01, ***p < 0.001). **C)** Intracellular calcium change measured using Fluo-4 AM in HEK293-hTREM2/DAP12 cells. Cells were treated with indicated concentrations of VG-3927 (red bar) or As48 (green bars) for 10 minutes. Calcium levels were normalized to vehicle control. Data are presented as mean ± SEM (n = 4-5). Statistical significance was determined by unpaired two-tailed t-test (**p<0.01, ***p < 0.001).

To evaluate downstream signaling, we measured PLCγ2-dependent calcium mobilization using the Fluo-4 AM calcium indicator. As48 induced a robust calcium response, with significant increases observed at concentrations as low as 1 μM, and up to a 5-fold elevation at 3–10 μM (Fig 5C).

Interestingly, As48 demonstrated greater potency in inducing calcium mobilization compared to SYK phosphorylation, indicating that distinct activation thresholds may exist across TREM2-dependent signaling pathways. Collectively, these findings establish As48 as a functional TREM2 agonist capable of activating both proximal (SYK phosphorylation) and distal (calcium signaling) components of the pathway, with a pharmacological profile that is distinct from that of VG-3927.

### As48 Activates TREM2 Signaling in Human Microglial Cells

To evaluate the functional activity of As48 in a physiologically relevant cell, we examined its ability to activate TREM2 signaling in human microglial cells. We first assessed the endogenous expression levels of TREM2 and its signaling adaptor DAP12 in HMC3 human microglial cells. Western blot analysis revealed that wild-type HMC3 cells express minimal levels of TREM2 protein, while DAP12 expression was readily detectable (Fig 6A). To enable assessment of TREM2 signaling activation, we generated HMC3 cells stably overexpressing human TREM2, which showed robust TREM2 protein expression while maintaining endogenous DAP12 levels. Using the HMC3 TREM2 overexpression (OE) cells, we investigated the temporal dynamics of TREM2 signaling activation by As48.

**Figure 6.**
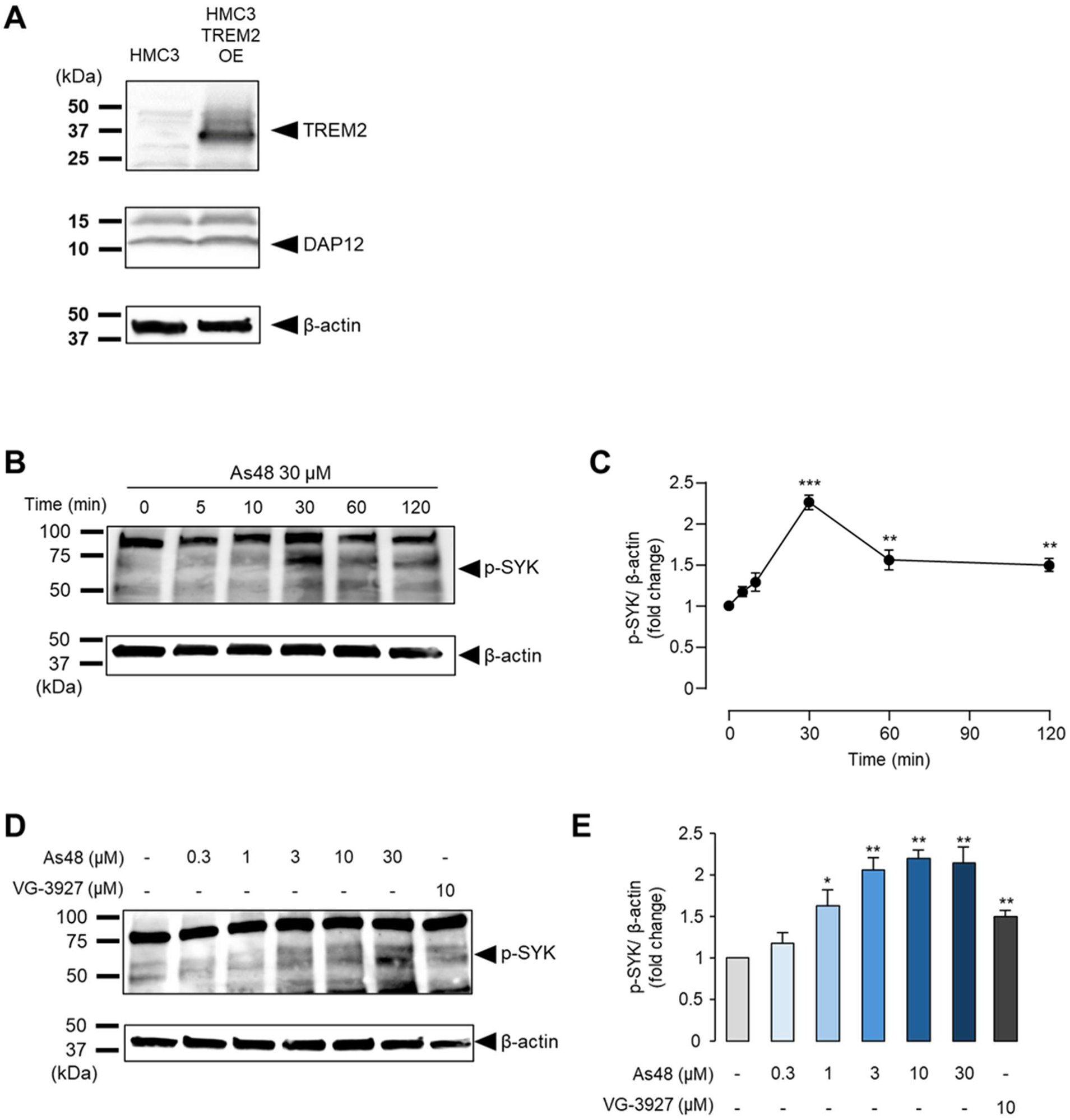
As48 activates TREM2 signaling in human microglial cells. **A)** Representative Western blot analysis showing TREM2 and DAP12 protein expression levels in wild-type HMC3 cells and HMC3 cells with TREM2 overexpression (OE). β-actin serves as loading control. **B,C)** Time-course analysis of SYK phosphorylation following As48 treatment (30 μM) in HMC3 TREM2 OE cells. **B)** Representative Western blots showing p-SYK levels at indicated time points (0-120 minutes). **C)** Quantification of p-SYK band intensity normalized to β-actin expression. Peak phosphorylation occurs at 30 minutes post-treatment. **D,E)** Dose dependent activation of SYK phosphorylation by As48 in HMC3 TREM2 OE cells. **D)** Representative Western blots showing p-SYK levels following 30-minute treatment with indicated As48 concentrations (0.3-30 μM). VG-3927 (10 μM) serves as positive control. **E)** Quantification of p-SYK band intensity normalized to β-actin expression, demonstrating dose-dependent TREM2 activation. Data are presented as mean ± SEM (n = 3). Statistical significance was determined by unpaired two-tailed t-test (*p < 0.05, **p < 0.01, ***p < 0.001).

Treatment with As48 (30 μM) induced rapid phosphorylation of SYK, a key downstream signaling mediator in the TREM2 pathway. Time-course analysis demonstrated that p-SYK levels peaked at 30 minutes post-treatment and gradually declined thereafter, reaching approximately 1.5-fold above baseline levels at 120 minutes (Fig 6B and C). This kinetic profile is consistent with typical TREM2 agonist activity and confirms that As48 can effectively trigger TREM2 signaling cascades. To determine the concentration-dependence of As48-mediated TREM2 activation, we treated HMC3 TREM2 OE cells with varying concentrations of As48 (0.3-30 μM) for 30 minutes.

Dose-response analysis revealed that As48 induced significant SYK phosphorylation starting at 1 μM, with maximal activation achieved at concentrations higher than 10 μM (Fig 6D and E). The potency of As48 was comparable to VG-3927 (10 μM), a well-characterized small-molecule TREM2 agonist used as positive control. These results demonstrate that As48 functions as an effective TREM2 agonist in human microglial cells, inducing dose-dependent activation of downstream signaling pathways with kinetics consistent with receptor engagement and activation. Although the experimental system required TREM2 overexpression due to low endogenous expression in HMC3 cells, these findings provide crucial evidence for As48 to activate TREM2 signaling in human microglial cells.

### As48 enhances microglial phagocytic function

Given that TREM2 activation promotes microglial phagocytosis—a critical neuroprotective mechanism in neurodegenerative diseases—we investigated whether As48 could augment phagocytic activity in BV2 murine microglial cells (Savage, Jay et al. 2015). Serum-deprived BV2 cells were treated with compounds for 30 minutes prior to exposure to fluorescent latex beads (30min), which served as phagocytic targets.

This two TREM2 agonists significantly increased phagocytic activity compared to vehicle control. VG-3927 (25 μM) increased the proportion of phagocytic cells by approximately 45%, whereas As48 (25 μM) induced a twofold increase (Fig 7A–D), despite As48 exhibiting weaker SYK phosphorylation than VG-3927 at equivalent concentrations. Both compounds enhanced phagocytosis, with As48 demonstrating a greater effect. These findings highlight the therapeutic potential of As48 in modulating microglial-mediated neuroprotection.

**Figure 7.**
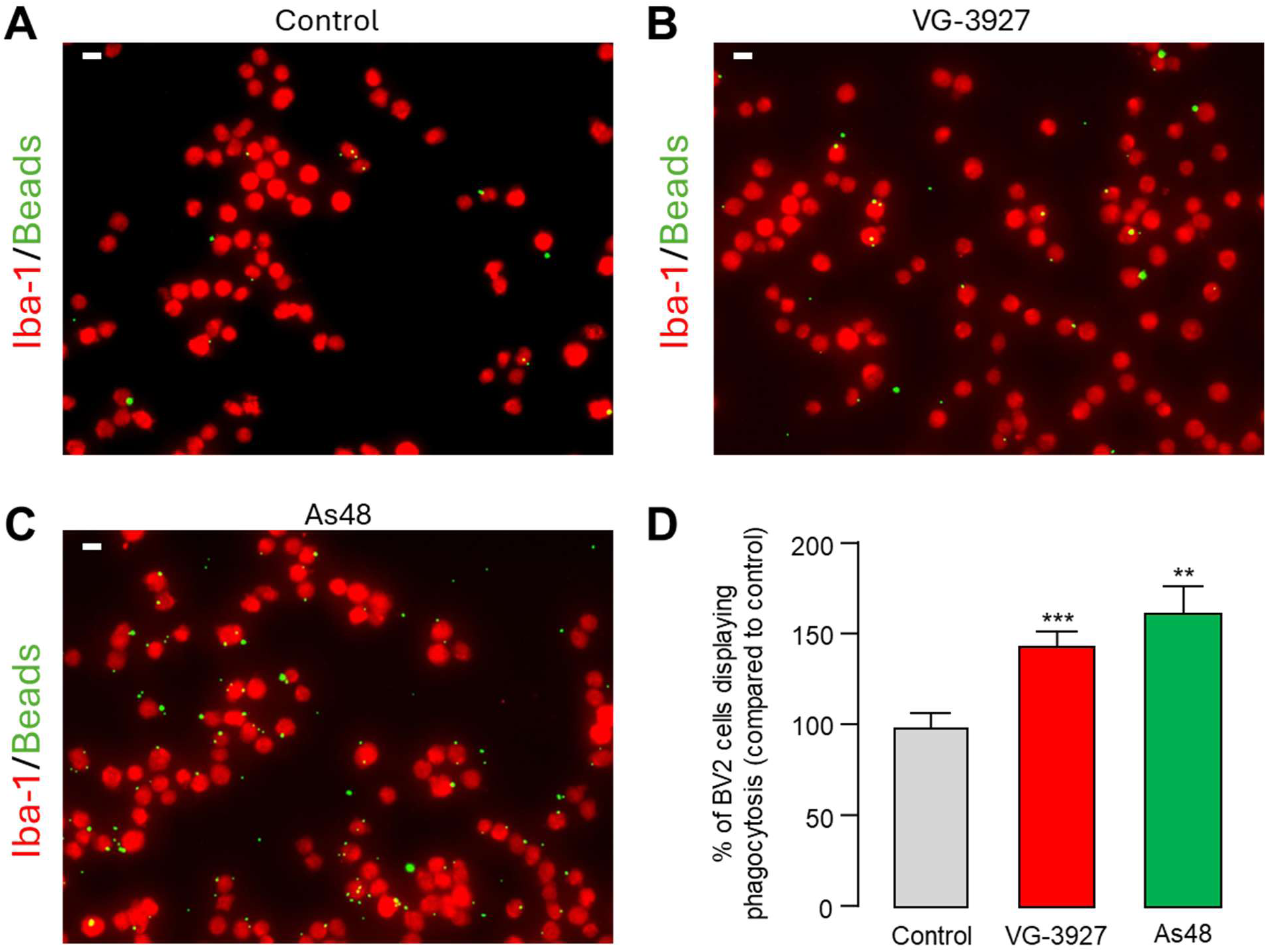
As48 enhances microglial phagocytic activity. **A-C)** Representative fluorescence microscopy images of BV2 murine microglial cells immunolabeled with anti-Iba-1 antibody (red) demonstrating microglial phenotype and containing fluorescent latex beads (green) internalized within the cell bodies. Cells were treated with **A)** Vehicle control (DMSO), **B)** VG-3927 (25 μM), and **C)** As48 (25 μM) treatment for 30 min. Scale bar = 10 μm. **D)** Quantification of phagocytic BV2 cells containing at least one internalized fluorescent bead, expressed as a percentage relative to vehicle control. Cells were treated with the indicated compounds (25 μM) for 30min prior to bead exposure (30min). Data are presented as mean ± SEM, based on analysis of 21–27 images per group from three independent experiments (N = 3). Data represent mean ± SEM from 3 independent experiments. Statistical significance was assessed using Brown-Forsythe and Welch Anova test followed by Dunnett’s multiple comparison test (*p<0.05, **p < 0.01, ***p < 0.001).

### Molecular Docking and Dynamics Simulation Reveal As48 Binding Mechanism

Based on the experimental observation of As48 binding to TREM2, docking and molecular dynamic simulations were carried out to investigate the binding mechanism of As48 with TREM2. Given the inherent flexibility of TREM2, the Induced Fit Docking (IFD) protocol in Maestro was employed to accurately account for conformational adjustments upon ligand binding. The molecular docking assay showed the presence of a well-defined binding pocket formed primarily by residues TYR61, Tyr62, Gly68, Ser138, and Leu233 (Fig 8C) which conformed with the observed experimental results. As48 established a hydrogen bond with Gly68 and several hydrophobic interactions which served to further stabilize the binding of As48 within the binding site.

**Figure 8.**
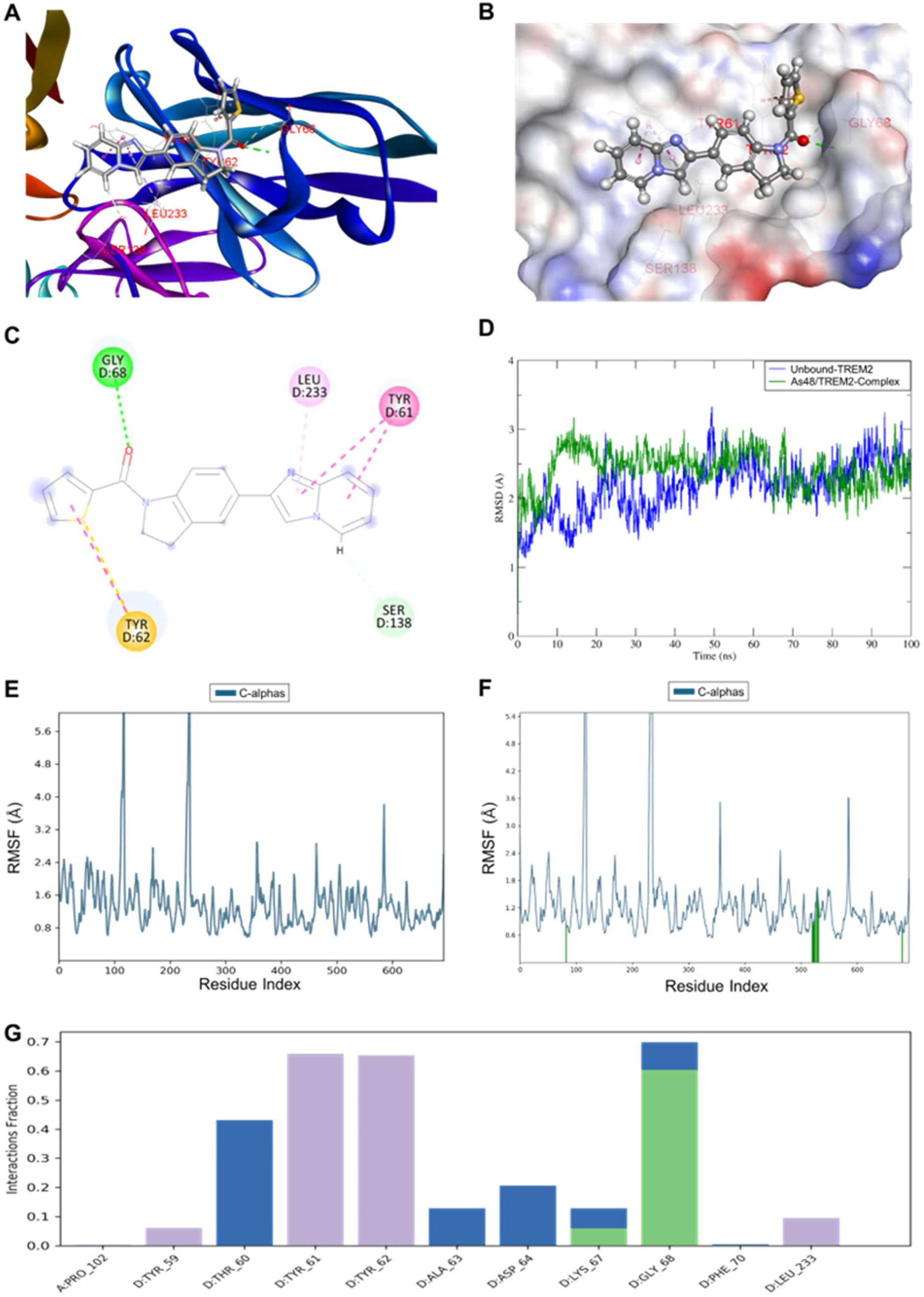
Molecular docking and MD simulation analysis of As48 binding to the His-tag region of TREM2. **A)** Docking pose of As48 within the cleavage site proximal region of TREM2, showing interactions with residues Tyr61, Tyr62, Gly68, Ser138, and Leu233. **B)** Surface representation of the TREM2 binding pocket with As48 positioned in the cleft, illustrating spatial complementarity and hydrogen bonding interactions. **C)** 2D interaction map of As48 with TREM2 residues, highlighting hydrogen bonds (green) and hydrophobic interactions. **D)** RMSD plot comparing backbone fluctuations of unbound TREM2 (blue) and the TREM2– As48 complex (green) over a 100 ns MD simulation, indicating overall structural stabilization upon ligand binding. **E-F)** RMSF plots showing residue-wise flexibility of TREM2 in the absence **E)** and presence **F)** of As48, with significant reduction in the fluctuations of the interacting residues in the bound state (green-highlighted regions). **G)** TREM2/As48 Interaction fraction analysis over the 100 ns of the MD simulation.

The RMSD analysis comparing the unbound and As48-bound TREM2 reveals that As48 binding stabilizes the global conformation of the receptor over 100 ns of simulation. The complex maintains a relatively stable backbone RMSD trajectory (blue, Fig 8D) which indicates that As48 does not introduce significant conformational disruptions but may reinforce local rigidity. This hypothesis is further supported by the RMSF comparison of the unbound and TREM2/As48 bound complexes behavior during the MD simulations. The unbound TREM2 (Fig 8E) displays high flexibility in regions around residue 59 to 138. In contrast, As48 binding (Fig 8F) reduces fluctuations in specific segments (particularly those around residues 58, 79 to 102) which correspond to direct interaction sites. These observations suggest that As48 binding restricts the mobility of flexible loop regions, which may potentially interfere with access or recognition by proteolytic machinery such as ADAM proteases.

The protein interaction bar chart (Fig 8G) shows that Gly68, Tyr61, and Tyr62 maintained contact with As48 during the majority of the MD simulation. Based on these observations, particularly the hydrogen bond with Gly68, As48 appears to stabilize within the binding site near the cleavage region. Such interaction persistence suggests the possibility that stabilization of this region may impair conformational changes or protease accessibility required for ectodomain shedding. Altogether, these findings provide structural and dynamic insight into how As48, by binding near the cleavage site region, may potentially exert inhibitory effects on TREM2 cleavage. By anchoring key residues involved in structural flexibility and protease recognition, As48 is predicted to potentially prevent conformational changes or restrict access to the cleavage site, which offers a mechanistic rationale for its possible anti-shedding activity.

### As48 Inhibits TREM2 Shedding Through a Novel Mechanism Independent of ADAM Protease Activity

Based on our molecular docking results suggesting that As48 binds near the TREM2 cleavage region, we investigated its potential to inhibit TREM2 ectodomain shedding. TREM2 undergoes constitutive proteolytic cleavage by ADAM10 and ADAM17 at the His157-Ser158 peptide bond, releasing the extracellular domain and leaving a membrane-bound C-terminal fragment. To assess As48’s effect on this process, we employed a dual-antibody approach using antibodies specific for the TREM2 extracellular domain (residues 19-134) and cytoplasmic domain (near Leu221). HEK293 cells overexpressing TREM2 were treated with varying concentrations of As48 for 18 hours, and TREM2 shedding was analyzed by Western blot. GM6001, a broad-spectrum metalloproteinase inhibitor known to block TREM2 shedding, served as a positive control. While the cytoplasmic domain levels remained relatively constant across all treatment conditions, As48 demonstrated a dose-dependent preservation of the extracellular domain (Fig 9 A,B). Significant protection against shedding was observed at concentrations higher than 10 μM, with maximal effect achieved at 30 μM, the same concentration that effectively activates TREM2 signaling.

**Figure 9.**
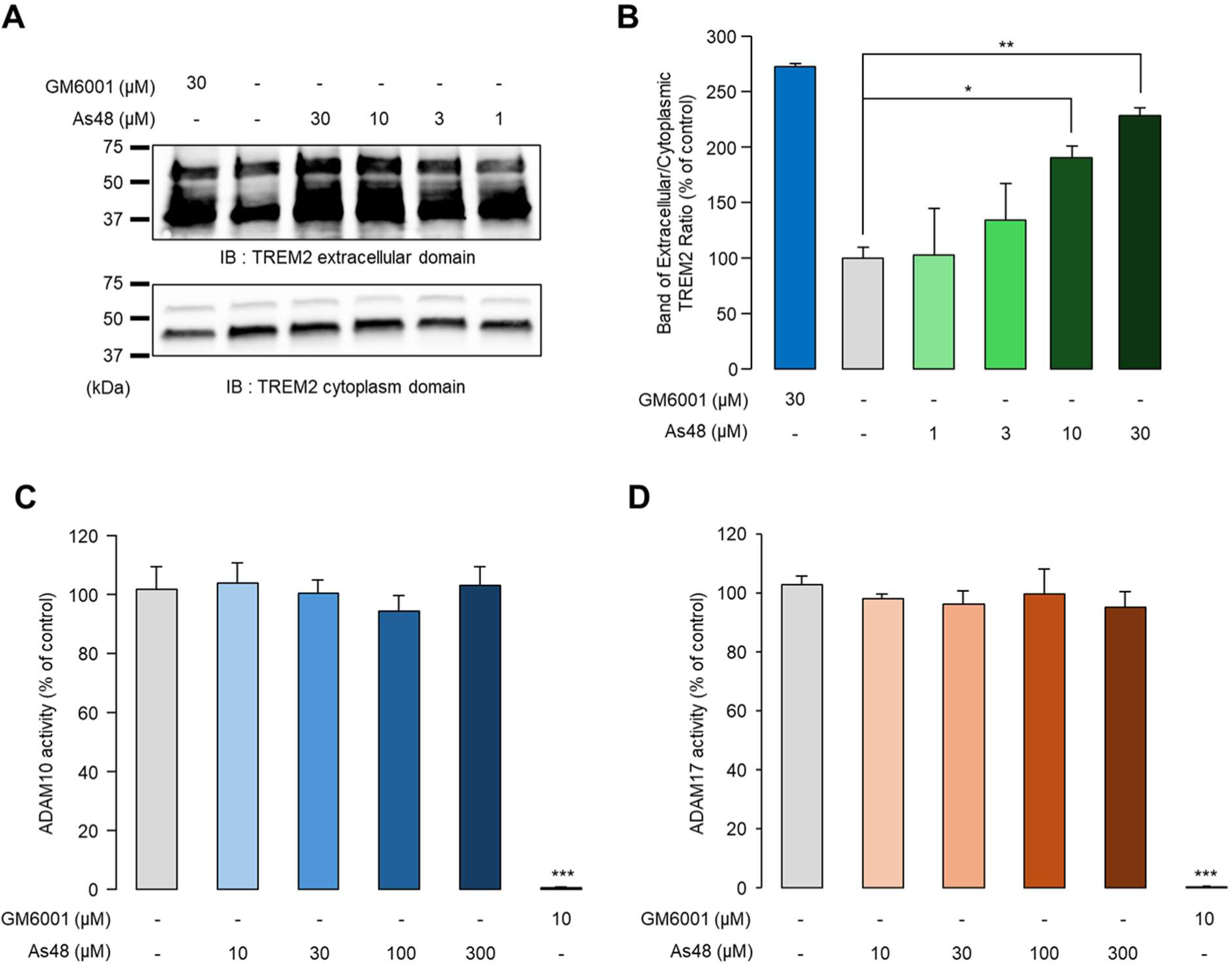
As48 inhibits TREM2 ectodomain shedding without affecting ADAM10/17 protease activity. **A)** Western blot analysis of TREM2 shedding in HEK293 cells overexpressing TREM2 following 18-hour treatment with indicated concentrations of As48. GM6001 (30 μM) and VG-3927 (10 μM) serve as positive and negative controls, respectively. Upper panel: TREM2 extracellular domain detection using antibody against residues 19-134; Lower panel: TREM2 cytoplasmic domain detection using antibody against the region near Leu221. **B)** Quantification of TREM2 shedding expressed as the ratio of extracellular domain to cytoplasmic domain band intensities. Mean ± SEM (n = 3). **C,D)** Assessment of ADAM10 **C)** and ADAM17 **D)** protease activities following As48 treatment at indicated concentrations. GM6001 (10 μM) serves as positive control for protease inhibition. Mean ± SEM (n = 3). Statistical significance was determined by unpaired two-tailed t-test (*p < 0.05, **p < 0.01, ***p < 0.001).

To determine whether As48 anti-shedding activity results from direct inhibition of ADAM proteases, we assessed the enzymatic activities of ADAM10 and ADAM17 using fluorogenic substrate assays. Unlike GM6001, which potently inhibited both ADAM10 and ADAM17 activities as expected, As48 showed no significant effect on either protease at concentrations up to 300 μM (Fig 9C and D). This finding indicates that As48 does not function as a direct ADAM protease inhibitor. These results demonstrate that As48 represents a novel class of TREM2 shedding inhibitor that operates through a mechanism distinct from direct protease inhibition. Consistent with our molecular docking predictions, As48 likely prevents TREM2 cleavage by inducing conformational changes near the cleavage site that render the His157-Ser158 bond inaccessible to ADAM proteases. This unique dual functionality, simultaneously activating TREM2 signaling while preventing receptor degradation, positions As48 as a potentially superior therapeutic modality compared to conventional TREM2 agonists that may be subject to enhanced shedding upon receptor activation.

### PK profiling of As48

To assess the developability of As48 as a CNS drug candidate for AD, we performed a comprehensive in vitro ADME and PK profiling campaign (Table 1). The compound displayed a balanced physicochemical profile, with a moderate logD7.4 of 2.4 and kinetic solubility of 75 µM in PBS (1% DMSO). The solubility of As48 in fasted state simulated intestinal fluid (FaSSIF) at 110 µM further supported oral absorption potential. Importantly, the BBB permeability of As48 was strongly favorable. In parallel artificial membrane permeability assay (PAMPA)–BBB, the compound exhibited high passive permeability (*Pₑ* = 7.1 × 10⁻⁶ cm/s), while in the MDCK–MDR1 assay it demonstrated high A→B transport (*P_app_* = 2.30 × 10⁻⁵ cm/s) and a low efflux ratio (ER = 1.10), suggesting negligible P-gp liability. Taken together, the PAMPA–BBB and MDCK–MDR1 data indicate that the CNS exposure of As48 will be governed predominantly by passive diffusion rather than transporter-mediated efflux, a profile predictive of efficient brain penetration.

**Table 1.** In vitro PK profile of As48.

| PK parameter | Value | PK parameter | Value |
| --- | --- | --- | --- |
| LogD<sub>7.4</sub><sup>a</sup> | 2.4 | t<sub>1/2</sub> [min] mouse plasma | 181 |
| Kinetic Solubility 1% DMSO/PBS [μM]<sup>b</sup> | 75 | t<sub>1/2</sub> [min] human plasma | 242 |
| Solubility in FaSSIF [μM] | 110 | t<sub>1/2</sub> [min] PBS | 95.1 ± 4.3 |
| PAMPA-BBB *P<sub>e</sub>* [cm/s] | 7.1 × 10<sup>-6</sup> | PPB human [%]<sup>c</sup> | 96.2 ± 0.24 |
| MDCK–MDR1 *P<sub>app</sub>* (A→B) | 2.30 × 10<sup>-5</sup> | IC<sub>50</sub> against WI-38 [μM] | >100 |
| MDCK–MDR1 *P<sub>app</sub>* (B→A) | 2.53 × 10<sup>-5</sup> | IC<sub>50</sub> against HS 27 [μM] | >100 |
| MDCK–MDR1 Efflux Ratio (B→A/A→B) | 1.10 | IC<sub>50</sub> against hERG [μM] | >50 |
| Stability in simulated gastric fluid (pH 1.2, 2h) | 92% remaining | CYP3A4 inhibition (% at 10 mM) | 12.7% |
| Stability in simulated intestinal fluid (pH 6.8, 2h) | 90% remaining | CYP2D6 inhibition (% at 10 mM) | 9.1% |
| t<sub>1/2</sub> [min] mouse liver microsomes<sup>c</sup> / CL<sub>int</sub><sup>d</sup> | 55.2 ± 3.2 / 25.1 ± 1.4 | CYP2C9 inhibition (% at 10 mM) | 7.4% |
| t<sub>1/2</sub> [min] human liver microsomes<sup>c</sup> / CL<sub>int</sub><sup>d</sup> | 70.8 ± 5.7 / 20.4 ± 2.3 | CYP2C19 inhibition (% at 10 mM) | 6.2% |

<sup>a</sup>LogD (pH = 7.4) the distribution coefficient at physiological pH 7.4. <sup>b</sup>Kinetic solubility measured in 1% DMSO/PBS (pH 7.4) by UV–vis spectrophotometry (n=3). <sup>c</sup>Values are mean ± SD (n=3).
<sup>d</sup>Intrinsic clearance expressed in μL/(min × mg) protein. Values are mean ± SD (n=3).

The compound also showed excellent stability in simulated gastric and intestinal fluids (>90% parent remaining after 2 h), as well as long plasma half-lives (181 min in mouse, 242 min in human). As shown in Table 1, the chemical stability of As48 in PBS was similarly robust (95.1 ± 4.3 min).

Metabolic turnover in liver microsomes was moderate, with half-lives of 55.2 ± 3.2 min (mouse, CL_int_ = 25.1 ± 1.4 μL/min/mg protein) and 70.8 ± 5.7 min (human, CL_int_ = 20.4 ± 2.3 μL/min/mg protein), indicating an acceptable clearance profile for a CNS drug candidate. Safety pharmacology assays showed no cytotoxicity in human fibroblasts (WI-38 and HS-27; IC₅₀ >100 µM), low hERG liability (IC₅₀ >50 µM), and minimal inhibition of major CYP isoforms (<13% at 10 µM), indicating a low risk of drug–drug interactions (Table 1). Overall, the ADME/PK results support the advancement of As48, demonstrating a favorable balance of solubility, BBB permeability, stability, and safety that are consistent with progression toward in preclinical testing in animal models of AD.

## Discussion

The identification of As48 represents a paradigm-shifting advancement in TREM2 pharmacology, demonstrating for the first time that a single small molecule can simultaneously activate TREM2 signaling while preventing receptor degradation through ADAM-mediated shedding. This breakthrough discovery was enabled by our innovative application of Affinity Selection-Mass Spectrometry (AS-MS) screening, which provided an unbiased, direct binding-based approach to TREM2 modulator discovery. Unlike conventional functional screening approaches that may miss compounds with unique mechanisms or those masked by TREM2 activation-induced shedding effects, AS-MS enabled the identification of 62 initial candidate molecules that directly bind to TREM2 (Dueñas, Peltier-Heap et al. 2023, Zhang, Calvo-Barreiro et al. 2024). Through orthogonal biophysical validation using TRIC, MST, and SPR, we successfully identified As48 as a micromolar-affinity TREM2 binder (K_D_ = 13.8 ± 1.08 μM), revealing its unprecedented dual functionality that combines receptor activation with shedding protection in a single molecule.

The most remarkable feature of As48 is this dual functionality that distinguishes it from existing TREM2 modulators. While monoclonal antibodies suffer from restricted blood-brain barrier penetration and manufacturing complexity (Pardridge 2012, Schlepckow, Monroe et al. 2020), and small molecule agonists like VG-3927 cannot address constitutive receptor shedding that depletes functional TREM2 from the cell surface (Kleinberger, Yamanishi et al. 2014, Feuerbach, Schindler et al. 2017, Mirescu 2024) As48 overcomes these limitations through its unprecedented mechanism. Our molecular dynamics simulations revealed that As48 binds to a well-defined pocket formed by residues TYR61, Tyr62, Gly68, Ser138, and Leu233, leading to stabilization of the broader flexible region (residues 59-138) with particular reduction in fluctuations around residues 58 and 79-102, which may restrict protease accessibility to the His157-Ser158 cleavage site (Heslegrave, Heywood et al. 2016). Importantly, this anti-shedding effect occurs without directly affecting ADAM10/17 enzymatic activity.

The differential potency observed between SYK phosphorylation (EC_50_ ∼10 μM) and calcium signaling (threshold ∼1 μM) provides important mechanistic insights, suggesting that calcium signaling represents a more sensitive readout of TREM2 engagement due to signal amplification in the phospholipase C pathway (Peng, Malhotra et al. 2010). Supporting this, As48 demonstrated superior phagocytic enhancement compared to VG-3927 in BV2 cells despite exhibiting weaker p-SYK signaling, suggesting that the shedding-blocking effect contributes significantly to TREM2-mediated phagocytic function. This dissociation between proximal signaling strength and functional outcome underscores the therapeutic advantage of combining receptor activation with protection from degradation.

The 7-fold selectivity of As48 for TREM2 over TREM1 represents a crucial safety advantage, ensuring selective enhancement of beneficial microglial functions without triggering detrimental inflammatory cascades through TREM1 activation (Bouchon, Facchetti et al. 2001, Singh, Rai et al. 2021). This selectivity profile positions As48 as a precision therapeutic tool for neurodegenerative diseases like Alzheimer’s disease, where TREM2 dysfunction contributes to impaired microglial clearance and inflammatory dysregulation (Guerreiro, Wojtas et al. 2013, Zhao, Wu et al. 2018). The dual functionality may achieve more robust and sustained therapeutic effects compared to conventional TREM2 agonists by simultaneously activating beneficial pathways while preventing receptor degradation.

As48’s comprehensive pharmacokinetic profiling reveals favorable blood-brain barrier permeability (PAMPA-BBB Pe = 7.1 × 10⁻⁶ cm/s), which exceeds the threshold for CNS penetration (>4.0 × 10⁻⁶ cm/s) established for brain-penetrant compounds (Di, Kerns et al. 2003, Mensch, Melis et al. 2010). The minimal P-glycoprotein efflux (efflux ratio = 1.10) indicates CNS exposure governed by passive diffusion rather than active efflux, further supporting brain bioavailability (Pardridge 2019). While the moderate metabolic stability [t1/2 = 55.2 min (mouse), 70.8 min (human)] and favorable safety profile support advancement toward preclinical evaluation, further optimization of both binding affinity and metabolic stability would enhance clinical development potential. The current binding affinity (KD 13.8 μM by MST, 29.5 μM by SPR) represents a starting point for medicinal chemistry optimization to achieve sub-micromolar potency typical of clinical candidates (Banks 2016).

However, several limitations of this study should be acknowledged. In vitro studies primarily utilized overexpression systems due to low endogenous TREM2 expression, necessitating validation in primary microglial cells or iPSC-derived microglia. The anti-shedding mechanism requires validation through direct measurement of soluble TREM2 levels in physiologically relevant systems, particularly in TREM2 variants with enhanced shedding such as the H157Y mutation (Kleinberger, Yamanishi et al. 2014, Schlepckow, Monroe et al. 2020). Additionally, evaluation in humanized animal models with direct measurement of cerebrospinal fluid sTREM2 concentration changes would provide crucial in vivo validation, as CSF sTREM2 levels serve as biomarkers for TREM2 activity in neurodegeneration (Suárez-Calvet, Kleinberger et al. 2016). The long-term consequences of sustained shedding inhibition on TREM2 biology and potential compensatory mechanisms require investigation in chronic dosing studies.

As48 represents a groundbreaking paradigm shift in TREM2 pharmacology by establishing the first small molecule capable of combining receptor activation with protection against degradation. This dual functionality addresses the fundamental limitation that has hindered previous TREM2 therapeutic approaches, where conventional agonists remain vulnerable to enhanced shedding upon activation. The discovery validates AS-MS screening as a powerful approach for identifying novel therapeutic modalities and establishes a new therapeutic paradigm that may inspire similar approaches for other challenging neuroinflammatory targets. Future studies should focus on affinity and metabolic stability optimization, validation in disease-relevant models, and comprehensive evaluation of the therapeutic window for dual TREM2 modulation. The identification of As48 not only provides a promising therapeutic candidate but also establishes proof-of-concept that dual-function TREM2 modulators represent a superior therapeutic strategy, opening new avenues for developing next-generation neuroinflammatory therapeutics that can simultaneously activate beneficial pathways while preventing receptor degradation.

## Acknowledgements

This work was supported by the National Institute on Aging under grant number R01AG083512 (PI: Gabr). Biophysical characterization experiments were performed using instrumentation at the Drug Discovery Research Center (DDRC) at The Rockefeller University, including the Dianthus instrument for TRIC, the Prometheus Panta system for thermal shift assays, and SPR instrumentation for binding kinetics analysis.

